# A fast and general method to empirically estimate the complexity of brain responses to transcranial and intracranial stimulations

**DOI:** 10.1101/445882

**Authors:** Renzo Comolatti, Andrea Pigorini, Silvia Casarotto, Matteo Fecchio, Guilherme Faria, Simone Sarasso, Mario Rosanova, Olivia Gosseries, Mélanie Boly, Olivier Bodart, Didier Ledoux, Jean-François Brichant, Lino Nobili, Steven Laureys, Giulio Tononi, Marcello Massimini, Adenauer G. Casali

## Abstract

**Background:** The Perturbational Complexity Index (PCI) was recently introduced to assess the capacity of thalamocortical circuits to engage in complex patterns of causal interactions. While showing high accuracy in detecting consciousness in brain injured patients, PCI depends on elaborate experimental setups and offline processing and has restricted applicability to other types of brain signals beyond transcranial magnetic stimulation and high-density EEG (TMS/hd-EEG) recordings.

**Objective:** We aim to address these limitations by introducing PCI^ST^, a fast method for estimating perturbational complexity of any given brain response signal.

**Methods:** PCI^ST^ is based on dimensionality reduction and state transitions (ST) quantification of evoked potentials. The index was validated on a large dataset of TMS/hd-EEG recordings obtained from 108 healthy subjects and 108 brain injured patients, and tested on sparse intracranial recordings (SEEG) of 9 patients undergoing intra-cerebral single-pulse electrical stimulation (SPES).

**Results:** When calculated on TMS/hd-EEG potentials, PCI^ST^ performed with the same accuracy as the original PCI, while improving on the previous method by being computed in less than a second and requiring a simpler set-up. In SPES/SEEG signals, the index was able to quantify a systematic reduction of intracerebral complexity during sleep, confirming the occurrence of state-dependent changes in the effective connectivity of thalamocortical circuits, as originally assessed through TMS/hd-EEG.

**Conclusions:** PCI^ST^ represents a fundamental advancement towards the implementation of a reliable and fast clinical tool for the bedside assessment of consciousness as well as a general measure to explore the neuronal mechanisms of loss/recovery of brain complexity across scales and models.

## Introduction

Measures of brain complexity have recently begun to move from the realm of theoretical neuroscience [1-6] into the field of experimental neurophysiology to study differences between global brain states, from wakefulness to sleep and anesthesia [7-10]. Further, measures of brain complexity have been considered as useful paraclinical indices to assess consciousness at the bedside of brain-injured patients [11-15]. In this spirit, a novel strategy based on quantifying the global effects of direct cortical perturbations was recently introduced [16]. This approach is motivated by the general theoretical principle that a brain’s capacity for consciousness relies on its ability to integrate information [5]. In this perspective, a critical mechanism supporting the emergence of conscious experience is the ability of different neural elements to engage in complex patterns of causal interactions such that the whole system generates information over and above its parts.

Practically, in order to estimate the amount of causal, irreducible information that a system can generate, a general procedure was implemented based on two steps: (i) locally perturbing the system in a controlled and reproducible way to trigger a cause-effect chain and (ii) quantifying the spatiotemporal complexity of the ensuing deterministic response to estimate information. The original implementation of this perturb-and-measure approach [16] involved (i) stimulating the brain with transcranial magnetic stimulation (TMS) and (ii) computing the algorithmic (Lempel-Ziv) complexity of the resulting patterns of activations at the level of cortical sources derived from the inverse solution of high-density electroencephalographic (hd-EEG) responses; this metric will be henceforth referred to as Lempel-Ziv Perturbational Complexity Index (PCI^LZ^).

Albeit macroscopic and coarse, PCI^LZ^ provided maximum (100%) accuracy in detecting consciousness in a large (n=150) benchmark population of subjects who could confirm the presence or absence of conscious experience through immediate or delayed reports [17]. PCI^LZ^ was lower in all unresponsive subjects who did not report any conscious experience upon awakening from NREM sleep or midazolam, xenon, and propofol anesthesia, and was invariably higher in conditions in which consciousness was present, including awake controls, conscious brain-damaged patients and subjects who were disconnected and unresponsive during dreaming and ketamine anesthesia but retrospectively reported having had vivid conscious experiences upon awakening [17, 18]. Once calibrated on the gold-standard of subjective reports, PCI^LZ^ measurements performed at the bedside of non-communicating subjects with brain injuries offered high sensitivity (94%) in detecting minimally conscious patients and allowed identifying a significant percentage (about 20%) of vegetative state/unresponsive wakefulness syndrome (VS/UWS) cases with high brain complexity, who had a higher chance of eventually recovering consciousness [17].

While PCI^LZ^ performs with unprecedented accuracy, it also has practical drawbacks and limitations. First, PCI^LZ^ can only be computed on spatiotemporal matrices of cortical activations that are obtained after an intensive processing of TMS/hd-EEG data, including forward modeling [19], source estimation [20] and permutation-based statistics at the single-trial level. All these steps imply a complicated and lengthy off-line analysis pipeline that hinders the dissemination of the method and its application as a routine clinical bedside tool. Clearly, the possibility of estimating perturbational complexity directly at the level of EEG sensors may have critical advantages: not only it would render the analysis process faster (ideally, on-line), easier to standardize and immune to the technical caveats of source modeling, but it would also allow the use of simplified and cheaper set-ups (i.e. not requiring hd-EEG and subject-specific MRI scans).

A second important drawback of PCI^LZ^ is its limited application to signals other than TMS/hd-EEG evoked potentials. Intracranial stimulations/recordings in humans [21, 22] and in animal models [23-26] as well as intra and extracellular responses recorded from cortical slices [27, 28] offer an unprecedented range of opportunities to interpret the TMS-EEG results and to elucidate the relationships between neuronal dynamics, network complexity and consciousness [29]. However, because PCI^LZ^ relies on EEG source estimation, its extension to other types of recordings, such as sparse matrices of intra-cerebral stereo EEG (SEEG) recordings and in vivo/in vitro local field potentials, is not straightforward [27].

Here we address these limitations and propose a novel measure of perturbational complexity that bears conceptual similarities with PCI^LZ^ but is much faster to compute and in principle generalizable to any type of brain signal evoked by perturbations or event related. Conceptually, we started from the notion that the binary sequences of activation and deactivations which are compressed by PCI^LZ^ can be considered as sequences of transitions between different states: a “response state” and a “non-response” or “baseline state” [30]. Thus, one should expect to find high values of perturbational complexity in systems that react to the initial perturbation by exhibiting multiple and irreducible patterns of transitions between response and non-response states.

Following this intuition, we developed PCI^ST^, an index that combines dimensionality reduction and a novel metric of recurrence quantification analysis (RQA) to empirically quantify perturbational complexity as the overall number of non-redundant state transitions (ST) caused by the perturbation. We validated PCI^ST^ on a large dataset of 719 TMS/hd-EEG sessions recorded from healthy subjects during wakefulness, non-rapid eye movement (NREM) sleep and anesthesia as well as brain-injured patients with disorders of consciousness (DOC). Finally, we tested PCI^ST^’s ability to probe the complexity of causal interactions within the brain by applying it on 84 SEEG recordings obtained in epileptic patients undergoing intra-cerebral single-pulse electrical stimulation (SPES) for clinical evaluation during both wakefulness and sleep.

## Material and methods

### Participants

#### Healthy subjects

The benchmark dataset consisted of 382 TMS/hd-EEG sessions reported in previous works [16, 17, 31]. Data were recorded from 108 healthy subjects (female, n = 63; age range = 18–80 years) in two conditions: (1) while they were unresponsive and did not provide any subjective report upon awakening (NREM sleep, n = 19; midazolam sedation at anesthetic concentrations, n = 6; anesthesia with xenon, n = 6; anesthesia with propofol, n = 6) and (2) while they were awake and able to provide an immediate subjective report (n = 103, including 32 subjects also recorded in the previously described unresponsive conditions). Protocols and informed consents were approved by the local ethical committees [16, 17, 31].

#### Brain-injured patients

TMS/hd-EEG data were also obtained in a population of 108 brain-injured patients (95 reported in a previous work [17]) with newly acquired data recorded following the same previously reported protocol [17]. Sixteen brain-injured patients were conscious and encompassed 5 individuals affected by locked-in syndrome (LIS) and 11 individuals who emerged from minimally conscious state (EMCS) by recovering functional communication and/or functional use of objects after a previous DOC. The remaining 92 brain-injured patients had a severe DOC and were repeatedly evaluated with the Coma Recovery Scale-Revised (CRS-R) for a period of 1 week (4 times, every other day). Patients showing only reflexive behavior across all evaluations were considered as being unresponsive (Unresponsive Wakefulness Syndrome, UWS, 43 patients), whereas patients showing signs of nonreflexive behaviors in at least one evaluation were considered as minimally conscious (Minimally Conscious State, MCS, 49 patients). Protocols and informed consents were approved by the local ethical committees [17] and written informed consent was obtained from healthy subjects, from communicative patients, and from legal surrogates of DOC patients.

#### Epileptic patients

Data included in the present study derived from a dataset collected during the pre-surgical evaluation of nine (eight previously reported [22]) neurosurgical patients with a history of drug-resistant, focal epilepsy. All subjects were candidates for surgical removal of the epileptic focus. During the pre-surgical evaluation all patients underwent individual investigation with SPES and simultaneous SEEG recordings for mapping eloquent areas and for precisely identifying the epileptogenic cortical network [32]. The investigated hemisphere, the duration of implantation and the location and number of stimulation sites were determined based on the non-invasive clinical assessment. The stimulation, recording and data treatment procedures were approved by the local ethical committee [22]. All patients provided written informed consent.

### TMS/hd-EEG measurements and data analysis

Specific protocols for acquiring and analyzing TMS/hd-EEG potentials were described in [16] and [17]. In brief, data were recorded with a 60-channel TMS-compatible EEG amplifier and MRI-guided TMS pulses were delivered with a focal biphasic stimulator. A noise masking sound tailored to the specific coil was played through inserted earphones and titrated by subjects within safety limits (<85dB). In each subject, multiple sessions of ∼200 stimuli were collected with TMS targeted to different areas at different intensities accordingly to the specific protocol. EEG responses to TMS were visually inspected to reject single trials and channels with bad signal quality. Independent component analysis was applied to remove residual artifactual components resulting from eye movements and muscle activations. Bad channels were then interpolated using spherical interpolation and data were bandpass filtered (0.1-45Hz), downsampled to 725 Hz, segmented between −400 and 400ms, re-referenced to the average, baseline corrected (−400 to −5 ms) and averaged across trials.

### Intracerebral measurements and data analysis

The procedures for SEEG data acquisition and analysis are described in [22]. Briefly, intracerebral activity was recorded at 1000Hz using a 192-channel recording system (Nihon-Kohden Neurofax-110) during electrical stimulation applied through one pair of adjacent contacts at different locations. SPES/SEEG sessions were obtained from all nine patients both during wakefulness and NREM sleep, resulting in a total of 84 sessions (see Table S1). Data were referenced to a contact located entirely in the white matter, subjected to linear detrend and bandpass filtering (0.5 – 300 Hz) and bipolar montages were calculated by subtracting the signals from adjacent contacts of the same depth-electrode [33, 34]. Stimulation artifact was reduced by applying a Tukey-windowed median filtering [35] between −5 and 5 ms. Data were segmented between −300 and 600ms and the SEEG magnitude at each electrode was computed as a z-score relative to its baseline. Trials and contacts showing pathological activity [36] were detected by visual inspection and excluded from the analysis and SPES-evoked responses were computed by averaging the remaining trials.

### Perturbational complexity index based on state transitions (PCI^ST^)

In its original formulation [16], perturbational complexity was calculated by binarizing TMS-evoked potentials (TEPs) at the EEG sources level using a fixed threshold derived from non-parametric statistics with respect to the baseline (pre-stimulus) and subsequently compressing the binary spatiotemporal patterns with the Lempel-Ziv algorithm. An underlying assumption of this strategy is that complex activations engaged by the perturbation appear on the evoked signals as patterns of oscillations around a fixed amplitude scale. Although proven successful when applied to the sources level, this approach was less sensitive in detecting complexity when calculated directly at the EEG-scalp level, where fast oscillations can appear riding on top of larger and slower envelopes as result of volume conduction and signal mixing (Figure S1). Furthermore, complex neuronal oscillations occurring in amplitude scales that are not determined by a fixed threshold can also be observed in microscopic and mesoscopic recordings due to cross-frequency couplings [37-41], and a binarized measure applied to such scales would have limited applicability to detect complex physiological activations.

Aiming at a general index of perturbational complexity that can be fast and efficiently calculated directly at the EEG-sensors level, we here took a non-binary approach and quantified the spatiotemporal complexity of evoked potentials by exploring multiple amplitude fluctuations present in principal components of the response. Starting from trial-averaged signals recorded in response to a perturbation, singular value decomposition was performed in order to effectively reduce the dimension of the data (Figure 1A). The principal components were selected so as to account for at least 99% of the response strength measured in terms of the square mean field power (see Supplemental Materials for definition and computational details) and components with low signal-to-noise ratio (SNR ≤ 1.1) were further removed.

**Figure 1.**
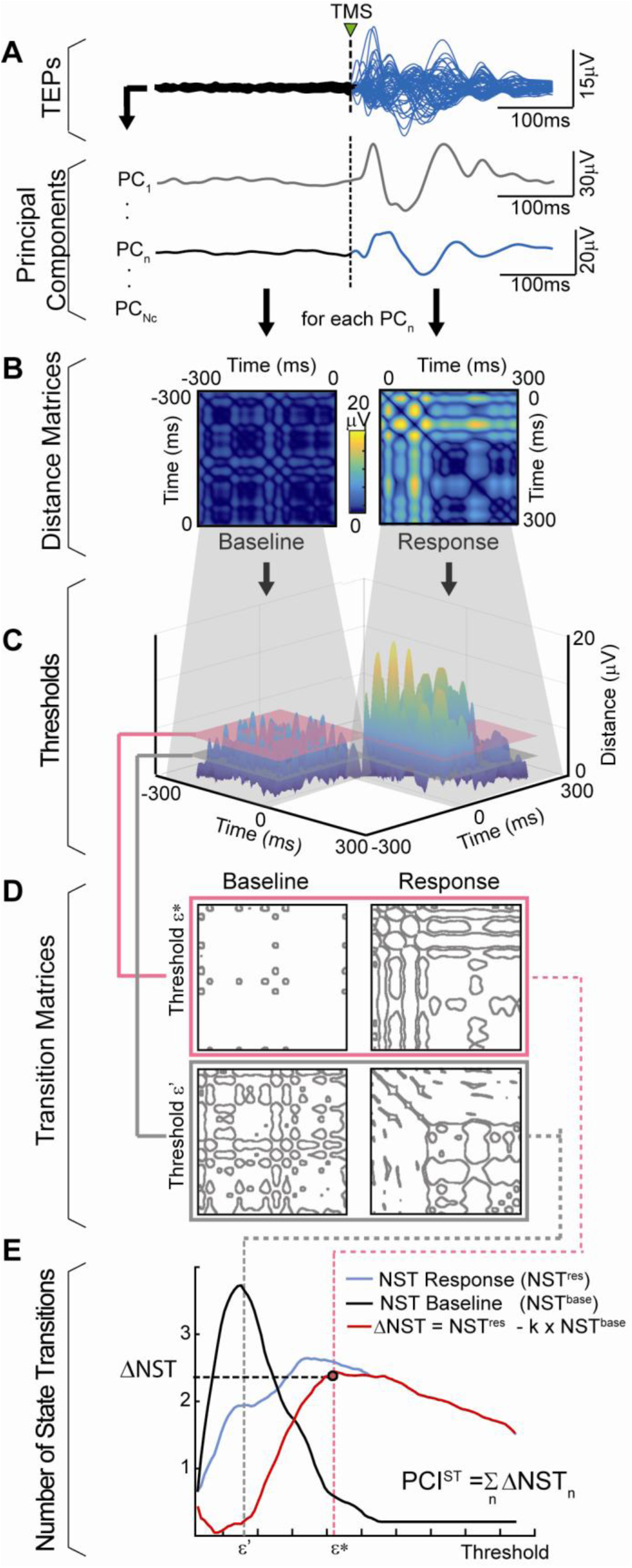
Calculating the Perturbational complexity index based on state transitions (PCI^ST^) from TMS/hd-EEG evoked potentials (TEP). PCI^ST^ is calculated by performing five steps: A) TEPs (butterfly plot, top) are decomposed in *N*_*c*_ principal components (PC) based on the singular value decomposition of the response to the perturbation. B) For each single component (PC_*n*_, highlighted) amplitude distances are calculated between every baseline samples (black trace in A) and between every response sample (blue trace in A), resulting in a baseline and a response distance matrix, respectively. C) These matrices are then thresholded at several scales. Two scale values are depicted in the figure: a lower threshold (*ε*’) and a higher threshold (*ε\**). D) At each scale, the corresponding transition matrices are computed for both baseline and response. These matrices are used to calculate the average number of state transitions (NST) in the baseline NST^*base*^ and in the response NST^*res*^. E) The complexity of the selected component is defined as the maximum weighted difference between the number of state transitions in the response and in the baseline (Δ NST_*n*_). The final measure PCI^ST^ is calculated by summing the Δ NST_*n*_ values across all *N*_*c*_ principal components.

The complexity of each resulting principal component was then evaluated using a method derived from RQA [30, 42, 43] by quantifying what we call *state transitions.* More specifically, distance matrices, defined by the voltage-amplitude distances between all time-points of the signal, were calculated separately for pre-stimulus and post-stimulus samples (Figure 1B). Next, these distance matrices were thresholded at a given scale *ε* (Figure 1C), yielding corresponding transition matrices (Figure 1D), i.e. contour plots that depicts the temporal transitions between states – roughly, the ups and downs in the signal –, for both the baseline and the response. By varying the threshold ε and comparing the average number of state transitions (NST) in the matrices of the response with that of the baseline, we looked for the scales at which the state transitions in the signal’s response were over and above the transitions present in the baseline activity. In this way, the complexity of the nth component (Δ NST_*n*_) was defined as the maximized weighted difference of NST between response (NST_*n*_^res^) and baseline (NST_*n*_^base^) signals:

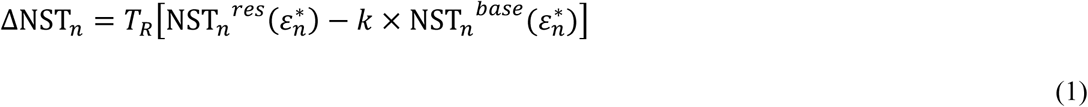

Where *ε*_*n*_^*^ is the threshold value which maximizes the weighted difference of NST (red dot in Figure 1E) and *T*_R_ is the number of samples in the response, a normalizing factor that yields a quantity that is extensive with the length of the signal’s response and largely independent of the sampling rate (Figure S2). The parameter *k* was introduced to control the relative weight between pre and post-stimulus state transitions. In this work, *k* was set to 1.2, value at which there is maximum separation between conscious and unconscious conditions (Figure S3).

Finally, PCI^ST^ was defined as the sum of these maximized significant state transitions across all principal components of the evoked perturbation signal:

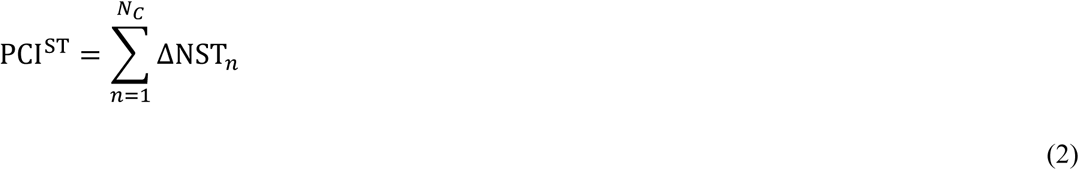

By its definition, PCI^ST^ is high when there are multiple linearly independent components in a spatially distributed response (spatial complexity), each one of them contributing with significant amounts of state transitions (temporal complexity) (Figure S4). Conversely, PCI^ST^ is expected to be low either if the perturbation evokes a strongly correlated response across different spatial recordings or if the independent components carry few temporal transitions in the response as compared to the baseline.

### Statistical Analysis

Data are presented as mean ± standard deviation (SD), and p-values less than 0.01 were considered significant. Wilcoxon-ranksum test was used for evaluating the discrimination between conscious and unconscious conditions. The classification power of discriminating different levels of consciousness was quantified by the area under the receiver operating characteristic (ROC) curve (AUC).

## Results and Discussion

### PCI^ST^ is reliable and fast in benchmark conditions

PCI^ST^ was calculated on a benchmark of 382 TEPs obtained in a group of 108 subjects during conscious (alert wakefulness) and unconscious (NREM sleep and anesthesia) conditions (Figure 2A). The wakefulness group presented significantly higher and more variable PCI^ST^ values (mean ± SD, 47.89 ± 12.65) than the NREM sleep/anesthesia group (14.19 ± 5.26, P = 4.7×10^−40^). In terms of classification performance between conscious and unconscious conditions, PCI^ST^ showed a high classification power that was equivalent to the performance of the original version of PCI in this same dataset [17] (AUC for PCI^ST^ =0.998, AUC for PCI^LZ^ = 0.995). Indeed, when values for each TMS/hd-EEG session were compared, we found a significant linear correlation between the metrics (r = 0.82, p<10^−95^, Figure 2B).

**Figure 2.**
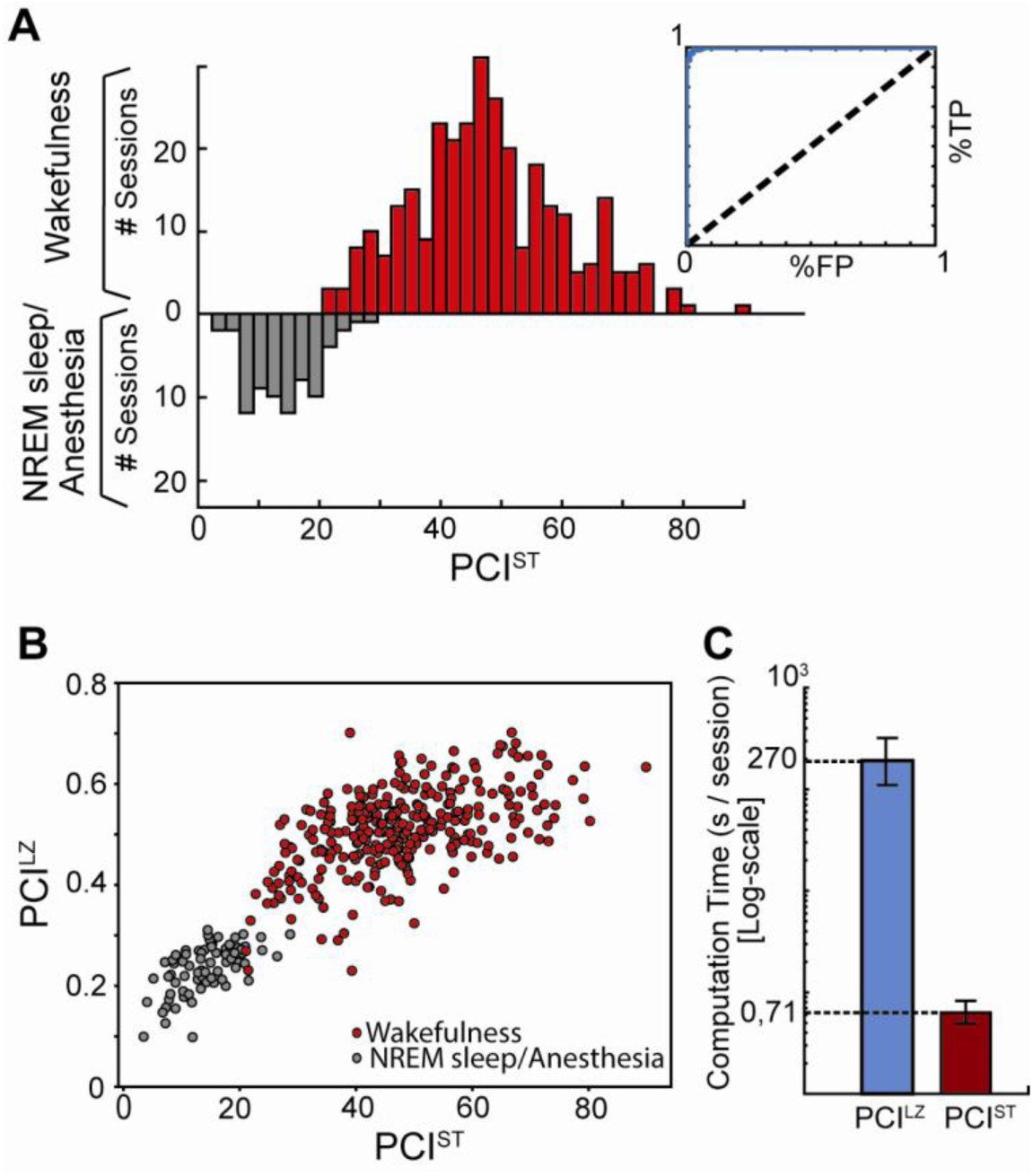
PCI^ST^ discriminates between consciousness and unconsciousness in healthy individuals and is faster than PCI^LZ^. (A) Histogram of PCI^ST^ values (left) for all 382 TMS sessions obtained from healthy individuals in the conscious (red) and unconscious (grey) conditions, with the corresponding ROC curve of the distributions (right). (B) Correlation between PCI^ST^ and PCI^LZ^ values in the benchmark dataset for conscious (red) and unconscious (grey) conditions (r=0.82, p<10^−95^). (C) Mean computation time per TMS/hd-EEG session for PCI^ST^ (red) and PCI^LZ^ (blue) calculated on the benchmark dataset.

On the other hand, because PCI^ST^ estimates perturbational complexity without employing source localization and surrogate techniques, PCI^ST^ computation was approximately 380 times faster than with PCI^LZ^. While PCI^LZ^ took about 300 seconds per session to compute (270s ± 99), PCI^ST^ was calculated in less than one second (0.71 ± 0.20, p< 10^−127^) (Figure 2C).

### PCI^ST^ allows a simple and fast set-up at the bedside of patients

We next tested the performance of PCI^ST^ in brain-injured patients. First, a threshold discriminating consciousness from unconsciousness was extracted from the PCI^ST^ values of the benchmark population using a linear classifier [44] (see Figure S5 for computational details). This empirical cutoff was then compared to PCI^ST^ values obtained from a group of 108 brain-injured patients who had recovered from coma and evolved toward various clinical conditions. Following the previous approach [17], we classified each patient using his maximum PCI^ST^ value obtained across all recorded sessions. This approach is aimed at assessing the patient’s best capacity for consciousness and parallels the diagnostic use of the best behavioral (CRS-R) score.

The sensitivity of PCI^ST^ in detecting signs of consciousness in brain-injured patients was comparable to PCI^LZ^ [17] (Figure 3A, top): PCI^ST^ made no erroneous classifications on conscious (LIS/EMCS) patients and achieved 91.9% sensitivity among minimally conscious individuals, correctly detecting signs of consciousness in 45 from 49 MCS patients (see Figure S6 for individual PCI^ST^ values).

**Figure 3.**
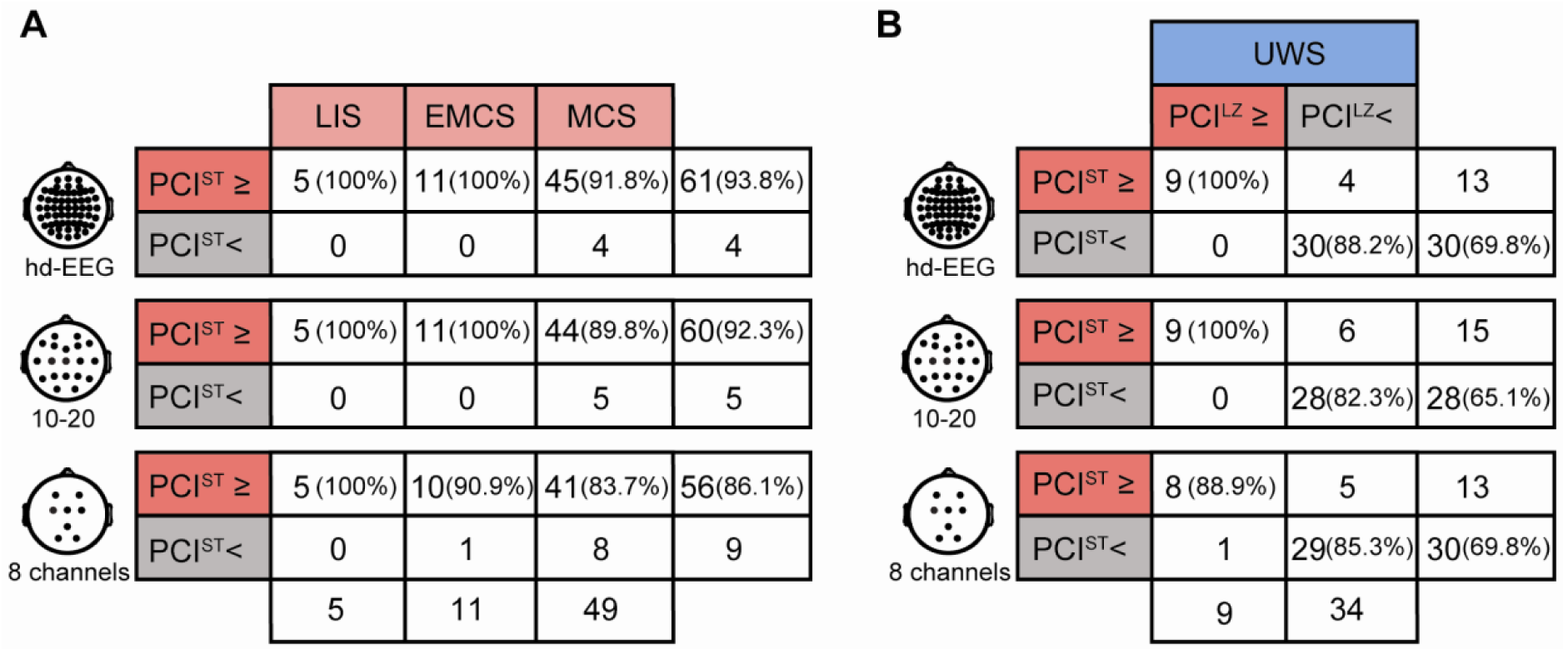
PCI^ST^’s ability to detect consciousness in brain-injured patients is preserved in simpler EEG set-ups. Number and percentages of patients classified as high (PCI^ST^≥) and low (PCI^ST^<) complexity with respect to the corresponding classification cutoffs obtained from the benchmark dataset are shown for EEG setups of 60 (top), 19 (middle) and 8 (bottom) channels. (A) PCI^ST^’ sensitivity in detecting signs of consciousness in conscious (EMCS/LIS) and minimally conscious (MCS) patients. (B) Contingency tables for the stratification of UWS patients in low complexity (PCI<) and high complexity (PCI≥) subgroups accordingly to PCI^LZ^ and PCI^ST^.

From a practical perspective, the potential of PCI to be employed as an index of consciousness in a clinical setting is significantly limited by the use of hd-EEG, which, besides entailing more expensive hardware, involves a cumbersome and lengthy preparation. While PCI^LZ^ necessarily demands hd-EEG systems so as to accurately perform source localization, PCI^ST^ can in principle be calculated on a reduced number of channels. We thus compared the performance of the index calculated on the original hd-EEG system (60 channels) to reduced setups containing 19 and 8 electrodes (see Supplemental Materials for further details). Notably the performance of the index diminished only slightly with the use of the standard 10-20 EEG system (19 channels), yielding sensitivities of 100% and 89.8% (44/49) for EMCS/LIS and MCS respectively (Figure 3A, middle). Finally, the simpler 8-channels setup resulted in reduced sensitivity scores on both EMCS/LIS (94%) and MCS (84%) patients (Figure 3A, bottom). An equivalent performance in discriminating conscious from unconscious conditions using simpler set-ups was also observed in the benchmark dataset (Figure S2).

In UWS patients, brain-based measures that do not require subject’s interaction with the external environment can be useful to detect a covert capacity for consciousness. In a previous study, PCI^LZ^ detected conscious-like complexity in 20.9% (9/43) of UWS patients, who also had a higher chance of recovery at 6 months [17]. Here, we evaluated whether these patients could also be identified by PCI^ST^. The novel index calculated on both high-density and standard 10-20 EEG setups detected all (n=9) the patients with high PCI^LZ^, whereas more than 82% of patients classified as low-complexity by PCI^LZ^ were also below threshold for PCI^ST^ (hd-EEG: 88.2%, 10-20 setup: 82.3%, Figure 3B-top and middle). The simpler 8-channels setup detected 8 out of 9 (88.9%) high-complexity patients and 29 out of 34 (85.3%) low-complexity patients (Figure 3B, bottom).

Taken together, these results show that PCI^ST^ calculation can afford an accurate and fast estimation of perturbational complexity even on reduced EEG set-ups. Combined with future optimizations of TMS-EEG hardware, this may allow the implementation of a practical, fast and potentially online method to be applied at the bedside in the routine clinical setting.

### PCI^ST^ reveals consistent changes of spatiotemporal complexity in intracerebral recordings

Beside its practical applications, estimating perturbational complexity based on state transitions enables the exploration of the brain’s causal structure across different recordings scales, from macroscopic EEG signals, to mesoscopic local field potentials and, in principle, to microscopic multisite electrophysiological/optical recordings. In the present study we explored this possibility at the mesoscale level by computing PCI^ST^ on sparse intracranial SPES-evoked potentials to test whether the state dependent changes in complexity revealed by TMS/EEG could be replicated and assessed by direct intracortical stimulation combined with SEEG recordings.

During wakefulness, the composite set of waves elicited by SPES appeared as a large number of components characterized by recurrent waves of activity lasting up to 600ms in the principal components space, which resulted in high PCI^ST^ values (Figure 4A-C). On the other hand, during NREM sleep, when SPES evoked a stereotypical wave, a small number of components were enough to span most of the response (Figure 4D-F). In addition, the few components that survived dimensionality reduction in NREM sleep showed fewer state transitions than the ones in wakefulness and accounted for a reduced PCI^ST^ value. These findings were reproducible across stimulation sites and consistent at population level (Figure 5). PCI^ST^ during NREM sleep was lower than in wakefulness for each one of the 42 different stimulation sites (Figure 5A-I). Overall, compared to wakefulness, PCI^ST^ was reduced during NREM sleep on average by 47.2% (Figure 5J) and significant at the group level (p = 1.4 × 10^−7^).

**Figure 4.**
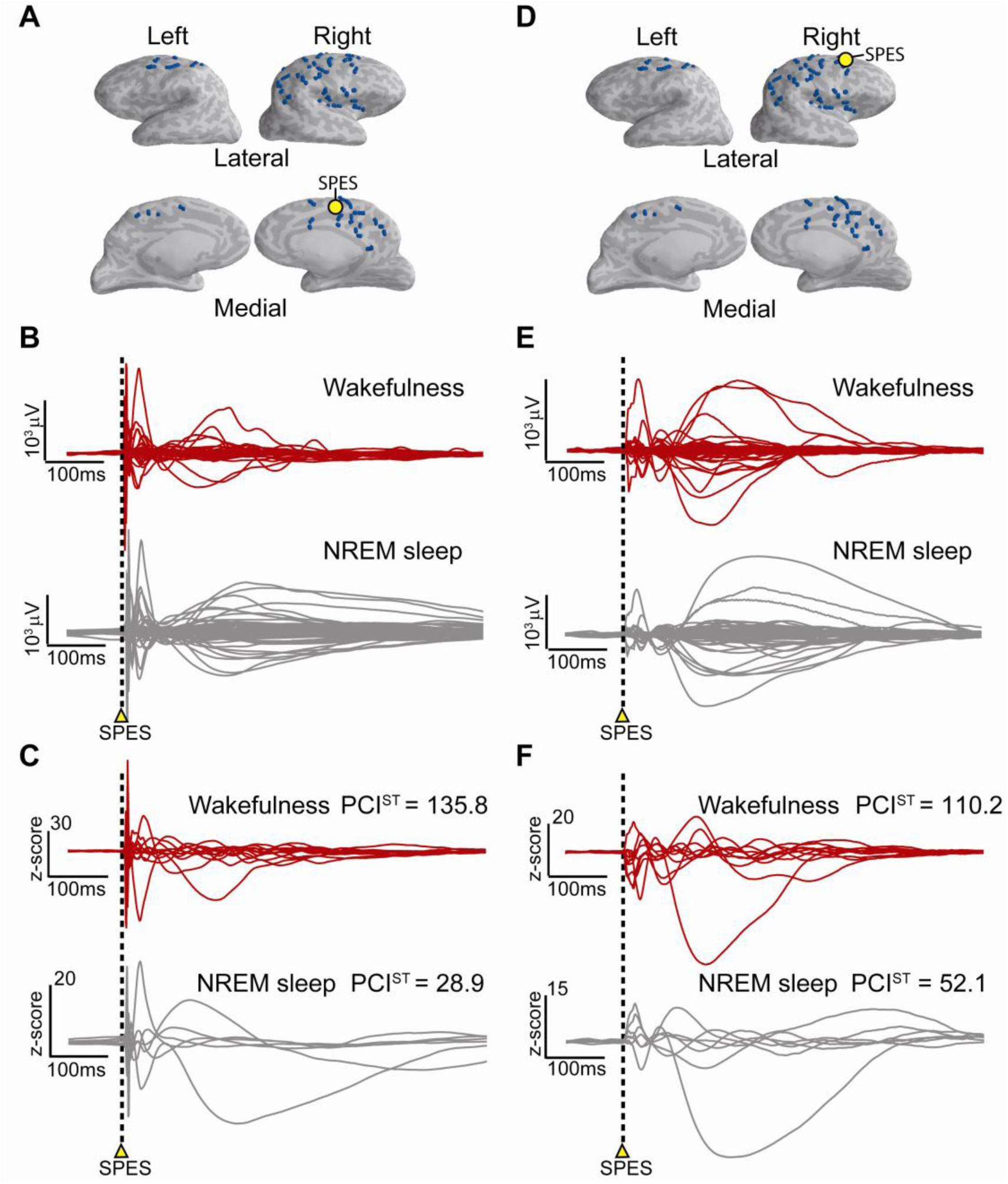
PCI^ST^ is able to quantify the spatiotemporal complexity of stereotactic EEG responses to SPES. SPES-evoked responses in the SEEG and principal component space are shown for a representative subject during stimulation delivered on the Superior Frontal Gyrus (panels A, B, C) and on the Superior Frontal Sulcus (panels D, E, F). Panels A and D depict the positions of the stimulating contact (yellow) and remaining SEEG contacts (blue) over a brain surface reconstructed from the individual’s brain. The correspondent SPES/SEEG-evoked responses are shown in the respective middle panels (B and E) as the superposition of the averaged SPES-evoked potentials recorded from all SEEG contacts during wakefulness (red traces) and NREM sleep (grey traces). Lower panels (C and F) depict the correspondent PCI^ST^ values and the normalized SPES-evoked responses decomposed in principal components after dimensionality reduction.

**Figure 5.**
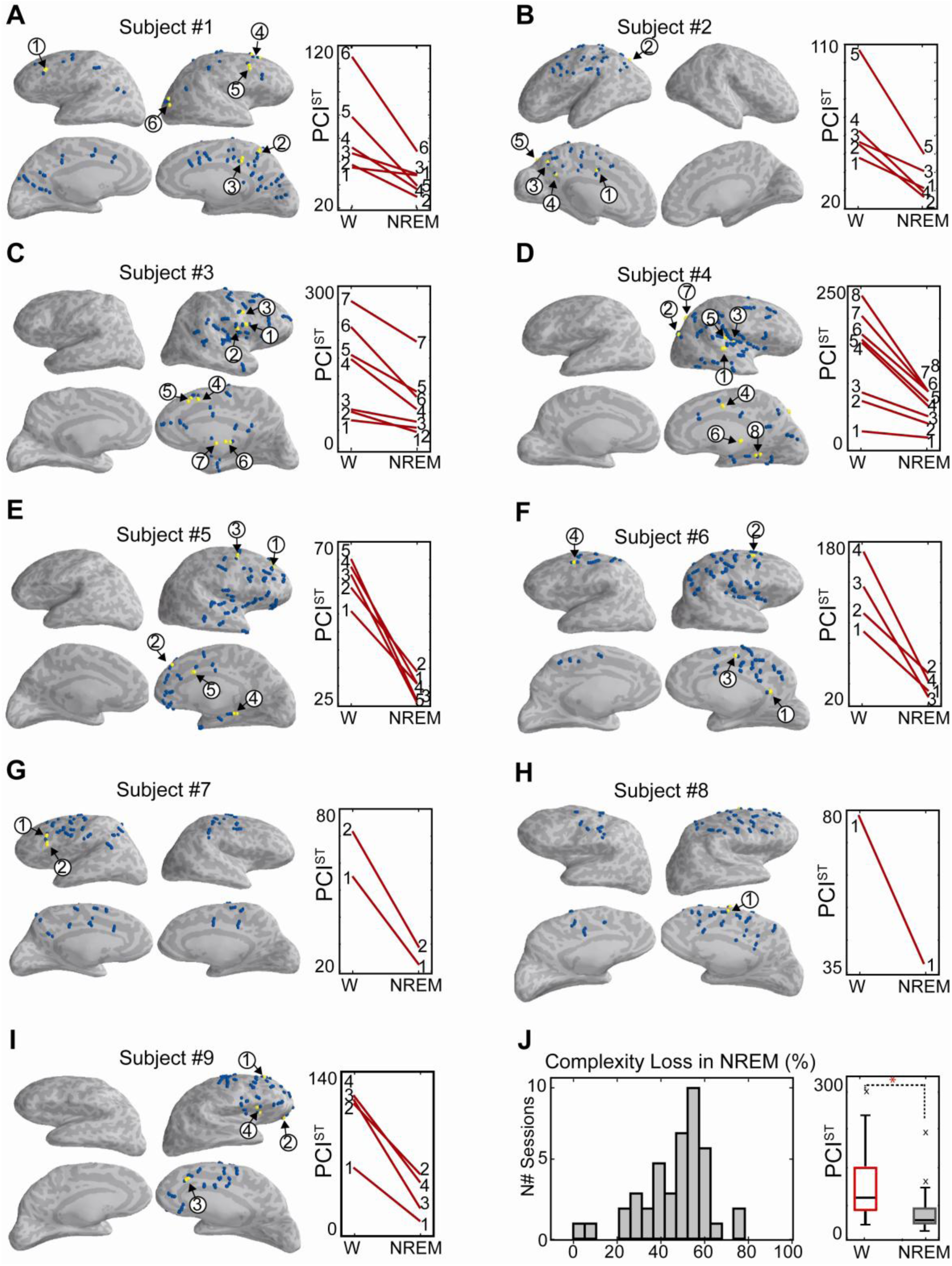
PCI^ST^ values in SPES-evoked potentials are invariably lower during NREM sleep as compared to wakefulness. Panels A – I: PCI^ST^ calculated in nine subjects during wakefulness (W) and NREM sleep are shown separately for each individual subject. SEEG (blue) and SPES contacts (yellow) are depicted over brain surfaces reconstructed from the individual’s brain (left). Numbers and arrows indicate the stimulation sites and the correspondent PCI^ST^ values (red traces, right). Panel J: shown are the percentage losses of complexity across all subjects and stimulation sites (left) and boxplots of PCI^ST^ values (right) at the group level for wakefulness (red box) and NREM sleep (grey box). Red asterisks indicate significant comparison (p = 1.4 × 10^−7^).

A key advantage of SPES over TMS is that the former is not associated with concurrent auditory and/or somatosensory stimulation [45]. The extent of the actual contribution of sensory co-stimulation to TEPs depends on many factors, such as coil type and effectiveness of noise masking, and is currently a matter of debate [46-49]. In this respect, the present intracranial results provide a definite confirmation of the fundamental interpretation of perturbational complexity, originally derived through TMS/hd-EEG recordings, as a genuine index of intracortical interactions. Direct intracortical stimulation elicited significant transitions contributing to the build-up of PCI^ST^ that occurred both at short and long latencies, that were specific for the stimulation site and state dependent (Figures 4 and 5). Thus, similarly to TMS, SPES elicited complex responses characterized by recurrent waves of activity in wakefulness and a stereotypical large-amplitude slow wave during NREM sleep. This finding is relevant as it confirms that the changes in perturbational complexity observed with TMS/EEG are not due to peripheral effects or subcortical sensory gating but reflect actual alterations in the intrinsic causal properties of thalamocortical circuits.

The application of PCI^ST^ also allows the exploration of the mechanisms of brain complexity at the finer scale of circuits and neuronal mechanisms. For example, at the level of the intracranial stimulation/recordings considered here, we observed substantial within-subject differences in the absolute values of PCI^ST^ depending on the stimulation site. These results suggest the presence of local differences in the ability of brain circuits to engage in complex patterns of causal interactions, which deserve further investigations.

Finally, the mesoscale assessment of perturbational complexity is in an ideal position to link microscale explorations at the bench to the macroscale measurements performed at the bedside of brain-injured patients. Connecting these levels is important as experiments in both cortical slices [27] and unresponsiveness wakefulness syndrome patients [31] suggest that loss of brain complexity is linked to the tendency of neurons to enter a silent period upon an initial activation (OFF-period). A thorough multiscale description of such mechanisms may be used to inform experimental and computational models [50] aimed at devising novel interventions to restore complexity and thus consciousness following brain injury.

## Conclusion

In this paper we have introduced, validated and tested PCI^ST^, a method of estimating perturbational complexity based on dimensionality reduction and state-transitions quantification. The novel index may not only provide a reliable, fast and potentially online option for the assessment of consciousness in the clinical setting, but also serve as a general translational tool for exploring the mechanisms of loss and recovery of brain complexity across species, scales, and models.

## Acknowledgments

We thank Nestor Caticha, Anna Cattani, Thierry Nieus, Simone Russo and Angela Comanducci for useful discussions during the course of this work. We thank Sasha D’Ambrosio for help with data collection and analysis.

