## Supplemental text, figures and table for "A fast and general method to empirically estimate the complexity of brain responses to transcranial and intracranial stimulations"

### Calculating PCI$^{ST}$

PCI combines a “perturb and measure” approach with a complexity metric in order to empirically estimate the amount of irreducible information generated through causal interactions within a system [1, 2]. This approach is unique because it permits, at least in principle, to explore the structure of causal interactions from the intrinsic perspective of the system under study, above and beyond the patterns of correlations that can be appreciated by a purely extrinsic, observational perspective [3, 4]. Further, by measuring the average response evoked by repeated stimulations of the same cortical target, only the effects causally related to the perturbation are amplified, whereas unrelated patterns of activity are averaged out. In this way, it is possible to explore effective interactions while minimizing the effect of noise, common drivers and elements that are causally segregated. Finally, to estimate the complexity of intrinsic causal interactions, only the irreducible (i.e. non-redundant) patterns of spatiotemporal significant activity are considered. Perturbational complexity is both spatial and temporal in nature inasmuch as it is only produced when different elements interact causally in an integrated and differentiated manner. In practice, perturbational complexity is high only when a perturbation propagates reliably over a set of integrated elements that react differently, giving rise to a transient response encompassing a variety of irreducible spatiotemporal patterns of activation.

In the case of PCI$^{ST}$, this transient response is interpreted as a trajectory in the multidimensional space spanned by the sensors. Mutanen et al. [5] has recently studied the ability of TMS to modulate brain activity by considering single-trial EEG responses to TMS as trajectories in the EEG signal space. Here we extended this geometrical approach and looked at the trajectories spanned by the evoked (trial-averaged) potentials. PCI$^{ST}$ estimates the spatiotemporal complexity of these evoked trajectories by performing two steps: dimensionality reduction and transition quantification.

#### Dimensionality reduction

In the first step, singular value decomposition (SVD) is used to rotate the trajectories into an orthonormal basis and optimally project them onto a space of lower dimension. This approach is similar to the concept of spatial PCA [6, 7], a data-driven technique that has been widely applied to evoked potentials [8-14] and that can be used to estimate the dimensionality of the trajectories in the EEG space in terms of the number of principal components accounting for the variance of the evoked response [15]. However, dimensionality reduction as operationalized in PCI$^{ST}$ differs from spatial PCA in two important ways: (i) PCI$^{ST}$ calculates SVD exclusively from the post-stimulus signals; (ii) the projection is based not on the covariance but on the correlation across sensors. This procedure guarantees that the principal components, onto which the evoked trajectories are projected, optimally explain the data in terms of the strength (mean field power) of the responses to the perturbation. More specifically, let **X** be the

More specifically, let **X** be the $N \times T$ matrix containing trial-averaged signals $x_n(t) = [x_n(t_1), \dots, x_n(t_k), \dots, x_n(t_T)]$ of a response to a perturbation recorded from $n = 1 \dots N$ different spatial locations and $k = 1 \dots T$ time samples. **X** can be represented by a trajectory $\mathbf{X}(t)$ in the $N$–dimensional space spanned by the spatial locations $\mathbf{e}_i$:

$$\mathbf{X}(t) = \sum_{i=1}^{N} x_i(t)\mathbf{e}_i$$

(1)

Let $t = 0$ be the instant of perturbation and $\mathbf{X}_{\boldsymbol{RES}}(t)$ ={ $\mathbf{X}(t) | t > 0\}$ be the response signal. Then let $\mathbf{a}_n$ be the normalized eigenvectors of the autocorrelation matrix **A** calculated from the response signal,

$$\mathbf{A} = \mathbf{X}_{RES} \times \mathbf{X}_{RES}{}^{T} = \left\{ \sum_{t>0} x_n(t) x_m(t) \,\middle|\, n = 1..N, m = 1..N \right\}$$

(2)

In the new orthogonal basis, $\{\mathbf{a}_n\}$, **X** can be represented by a rotated trajectory $\mathbf{Y}(t)$

$$\mathbf{Y}(t) = \sum_{n=1}^{N} y_n(t)\mathbf{a}_n = \sum_{n,m=1}^{N} x_m(t)(\mathbf{a}_n.\mathbf{e}_m)\mathbf{a}_n$$

(3)

The $N \times T$ matrix **Y** is thus composed by projecting the data **X** in the space spanned by the principal components (PC) obtained from SVD of the response signal. The PCs are then selected in terms of the eigenvalues of **A** sorted in decreasing order, so as to account for at least 99% of the response strength measured in terms of the square mean field power, $\sum_{n,k} |y_n(t_k)|^2$ . Next, components are selected in terms of their signal-to-noise ratio (SNR), which is calculated as the square root of the ratio of average response power (t>0, $T_R$ samples) to the average baseline power (t<0, $T_B$ samples):

$$SNR_n = \sqrt{\frac{\frac{1}{T_R}\sum_{t>0}|y_n(t)|^2}{\frac{1}{T_B}\sum_{t<0}|y_n(t)|^2}} \tag{4}$$

Components with low signal-to-noise ratio are removed, and the remaining $N_C$ responsive principal components are used for transition quantification (Figure 1A).

#### Transition quantification

While the number of selected principal components captures the spatial linear independence present in the response, the estimation of spatiotemporal complexity must also take into account the temporal information content present in each component. Since the complexity metric employed in the Lempel-Ziv formulation of perturbational complexity was based on binarizing temporal patterns with respect to a fixed significance threshold, $PCI^{LZ}$ shows a reduced performance when calculated in the absence of source modeling, such as in scalp EEG, when fluctuations of different frequencies relying on distinct neural mechanisms may appear linked due to volume conduction. In this case, complex patterns riding on top of larger and slower components may be easily missed (Figure S1). Crucially, sensitivity to cross-frequency patterns of fast oscillations that are modulated by slower envelopes are particularly important because they have been reported at multiple levels, from microscopic to macroscopic recordings [16-21], and because they may serve as the mechanism to coordinate neural dynamics and transfer information across spatial and temporal scales [22-25].

One way to explore the multiple amplitude scales of the activations in a time-series is by calculating its distance matrix [26, 27], which provides a summary of the geometrical relationships of the signal, not only in respect to the baseline but to every time point of the signal. Here, we proposed to quantify temporal complexity by binarizing not the original time-series but its corresponding distance matrix: a technique known as recurrence quantification analysis [27, 28].

We thus regard each principal component $y_n(t)$ separately and consider its time-delayed embedded form given by:

$$\mathbf{y}_n(t) = [y_n(t), y_n(t-\tau), \ldots, y_n(t - L \times \tau)]$$

(5)

where $L$ and $\tau$ are the dimension and the delay of the embedding, respectively and $n = 1 \ldots N_C$.. Given an embedded time-series $\mathbf{y}_n(t)$, its distance matrix $\mathbf{D}_n$ is then defined as the Euclidean distance between all timepoints of the signal:

$$\mathbf{D}_n(t_j, t_k) = \|\mathbf{y}_n(t_j) - \mathbf{y}_n(t_k)\|$$

(6)

Both matrices are then thresholded at several scales (Figure 1C): for each threshold $\varepsilon$ we calculate the transition matrix defined as the contour plot of the correspondent distance matrix at the scale $\varepsilon$ (Figure 1D):

$$\mathbf{T}_n(\varepsilon, t_j, t_k) = \begin{cases} 1, & if\ \mathbf{D}_n(t_j, t_k) = \varepsilon \\ 0, & otherwise \end{cases}$$

(7)

The transition matrix can be thought as the unidirectional derivative of the recurrence matrix $\mathbf{R}_n$, also known as recurrence plot [26]:

$$\mathbf{R}_n(\varepsilon, t_j, t_k) = \begin{cases} 1, & if\ \mathbf{D}_n(t_j, t_k) < \varepsilon \\ 0, & otherwise \end{cases}$$

(8)

Transition matrices can thus be interpreted as measuring not recurrent states, but transitions between states – between recurrent states, $\mathbf{R}_n(\varepsilon,t_i,t_j)$=1, and non-recurrent states, $\mathbf{R}_n(\varepsilon,t_i,t_j)$=0.

Recurrences are typically present in complex dynamical systems [29]: eventually every system will return to a state which is sufficiently close to a previous one and quantifying these recurrent states have been shown to be an effective way to detect rhythms and state changes within complex, possibly non-stationary, time-series [30-35]. Although inferring recurrences in the "true" state space of a dynamical system from a given time-series depends on properly choosing the embedding parameters to reconstruct the system's dynamical attractors [36], recurrence techniques are employed in calculating $PCI^{ST}$ simply to quantify patterns in the time-series, without assuming that the underlying attractors are faithfully reconstructed. Indeed, it has been shown that, when applied to experimental data, recurrence plots and the measures derived from them are able to quantify patterns in the dynamics even if no embedding is performed [37]. In the case of $PCI^{ST}$, independence on embedding dimension has been substantiated by the fact that the performance of the method in discriminating between brain states was found to be stable across different embedding parameters (Figure S3).

In the context of RQA, several metrics were proposed to quantify the complexity of the dynamics [27], such as the diagonal-line based entropy [35], the recurrence probability density entropy [38] or the Kolmogorov-Sinai entropy [39]. Although detecting recurrent states does not require signals to be stationary, these metrics are based on statistical properties of densities of recurrent states and may be unreliable in estimating complexity of very short and highly non-stationary time series, such as those evoked by a perturbation. We here took a different perspective and considered the number of transitions (NST) between recurrent and non-recurrent states in the principal components space as a measure of the information content of the response to a perturbation:

$$\mathrm{NST}_n(\varepsilon) = \frac{1}{{T_c}^2}\sum_{j=1}^{T_c}\sum_{k=1}^{T_c}\mathbf{T}_n\left(\varepsilon, t_j, t_k\right) \tag{9}$$

The NST quantifies the average amount of state transitions in a time-series across $T_C$ time samples at a given scale ε. Transitions are then averaged separately for both $T_C = T_B$ baseline samples (t<0), $\mathrm{NST}_n^{base}$ and $T_C = T_R$ response samples (t>0), $\mathrm{NST}_n^{res}$.

The threshold $\varepsilon$ represents a crucial amplitude scale through which information is encoded in the signal in terms of distance crossings. For very small scales (threshold $\varepsilon'$ in Figure 1D), most transitions occur in the baseline and the structures appearing in the Transition matrices at these scales are likely not caused by the perturbation. By increasing the value of $\varepsilon$, transitions start to appear mostly in the response to the stimulus and are, therefore, likely due to the perturbation. Thus, in order to measure the amount of temporal information encoded in the response over and above the amount encoded in the baseline, we search for the scale in which the weighted difference between $\mathrm{NST}_n^{res}$ and $\mathrm{NST}_n^{base}$ is maximized (threshold $\varepsilon^*$ in Figure 1D):

$$\varepsilon_n^* = \arg\max_{\varepsilon}\left[\mathrm{NST}_n^{res}(\varepsilon) - k \times \mathrm{NST}_n^{base}(\varepsilon)\right] \tag{10}$$

The parameter $k$ is introduced to control the relative weight between pre and post-stimulus state transitions. By setting the value of $k$, the crucial scale $\varepsilon_n$* becomes a data-driven parameter determined by the pre-post comparison for each principal component. Therefore, the temporal complexity of the $n$th component ($\Delta\mathrm{NST}_n$) is the maximized weighted difference at the optimal scale $\varepsilon_n$* scaled by the number of response samples ($T_R$) (Figure 1E). The scaling factor yields a quantity that is extensive with the length of the signal's response and largely independent on the sampling rate (Figure S2):

$$\Delta\mathrm{NST}_n = T_R\left[\mathrm{NST}_n{}^{res}(\varepsilon_n^*) - k \times \mathrm{NST}_n{}^{base}(\varepsilon_n^*)\right] \tag{11}$$

Finally, the State Transitions Perturbational Complexity Index is defined as the sum of the temporal complexity measured by $\Delta\mathrm{NST}_n$ taken across all the $N_\mathrm{C}$ selected principal components of the signal:

$$\mathrm{PCI}^{\mathrm{ST}} = \sum_{n=1}^{N_C} \Delta\mathrm{NST}_n \tag{12}$$

Notably, as this approach might be sensitive to patterns with a regular temporal structure that do not contribute to complexity as measured by compression-based metrics, $\mathrm{PCI}^{\mathrm{ST}}$ can be expected to accurately estimate complexity only when applied to transient short-lasting signals, such as evoked potentials, in which regular patterns repeating over a long time are unlikely to occur. Future studies in simulated dynamical systems [40, 41] and brain network models [42-45] should investigate to which extent counting state transitions with $\mathrm{PCI}^{\mathrm{ST}}$ is enough to capture the complexity of the causal structure of a system.

#### PCI$^{\mathrm{ST}}$ parameters

$\mathrm{PCI}^{\mathrm{ST}}$ was calculated on TMS/EEG signals and SPES/SEEG recordings. In both cases, a crucial step in calculating $\mathrm{PCI}^{\mathrm{ST}}$ is to control for stationary baseline-like activations that are likely not caused by the perturbation. This is done in two ways: (i) by removing principal components with low signal-to-noise ratio and (ii) by comparing between pre and post-stimulus transitions in the remaining components and selecting an amplitude scale that maximizes the information content of the response, over and above the content of the baseline. In this work, minimum SNR of the components accounting for 99% of the response strength was set to 1.1 and the *k* parameter controlling the relative weight between pre and post-stimulus state transitions in the remaining components was set to 1.2, value at which there is maximum separation between conscious and unconscious conditions (Figure S2). It is important to notice that these settings may be suboptimal for evoked signals with lower signal-to-noise ratio, such as those produced by peripheral stimulation, when it may be necessary to increase *k* and/or the minimum SNR to control for stationary baseline-like activations. Finally, since results were largely independent on the delay and embedding dimension (Figure S3), distance matrices of the principal components were calculated for the simplest case of L = 1.

$PCI^{ST}$ was calculated on TMS/EEG signals using three different EEG setups: hd-EEG (60 channels), standard 10-20 system with 19 channels (channels Fp1, Fp2, F7, F3, Fz, F4, F8, T3, C3, Cz, C4, T4, T7, P3, Pz, P4, T6, O1, O2) and a reduced setup with 8 channels (F3, F4, C3, Cz, C4, Pz, O1, O2). For each setup, signals were referenced to the corresponding common average reference.

Finally the baseline and response interval were respectively defined as (-400ms, -50ms) and (0ms, 300ms) for TMS/EEG sessions, and as (-300ms, -10ms) and (10ms, 600ms) for SPES/SEEG sessions – 0ms being the onset of the stimulation.

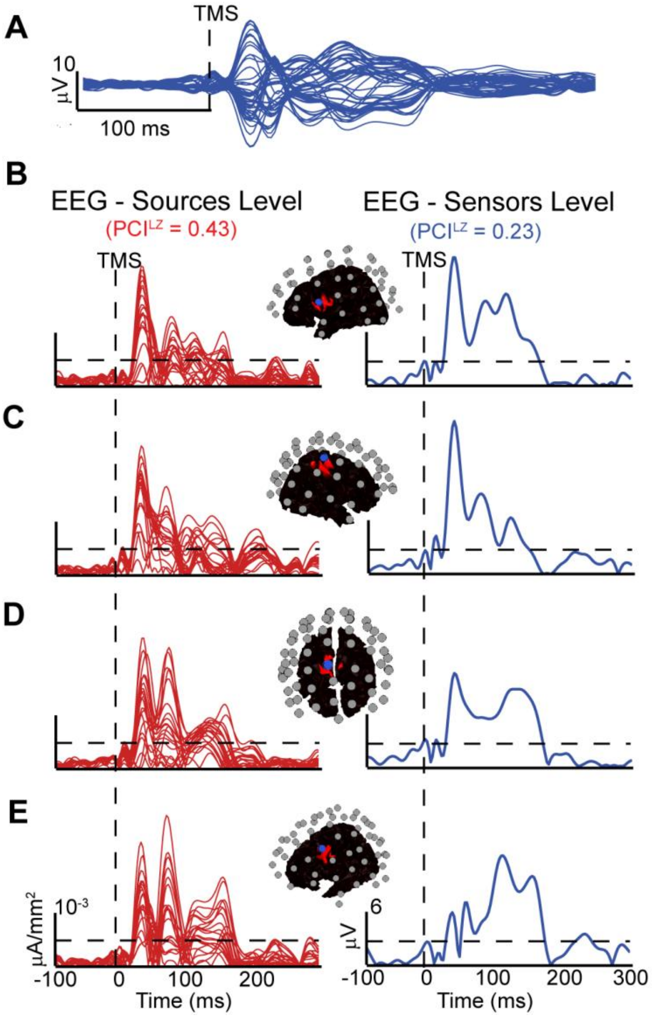


**Figure S1. Strategies based on binarizing TMS-evoked potentials across a fixed amplitude scale may fail to detect perturbational complexity at the scalp level**. (A) The blue traces show the superposition of the averaged TMS-evoked potentials (180 trials) recorded from all 60 EEG channels (butterfly plot) in a healthy subject during wakefulness. After source modeling, signals at the levels of cortical sources and scalp sensors were centralized on the mean and normalized on the standard deviation of their respective baselines (pre-stimulus). The time-series in panels (B) to (E) depict these normalized signals in absolute values: for each panel, the red traces (left) are the current densities for 20 dipolar sources located nearest to a particular EEG electrode, from which the corresponding voltage is displayed on the right (blue trace). Location of the depicted electrode (blue circle) and the corresponding dipolar sources (red patches) are shown on topographical maps of the cortex (black surface) together with the remaining electrodes (grey circles). At the source level (left), a fixed statistical threshold (horizontal dashed line), which was extracted from the distribution of amplitudes in the pre-stimulus, can be used to approximate the temporal complexity of the evoked response in terms of complex binary patterns of threshold crossings. At the scalp level (right), the fast deflections evoked by the perturbation are observed riding on top of slower and larger envelopes, thus causing methods based on the oscillations around a fixed amplitude scale to fail in detecting complexity.

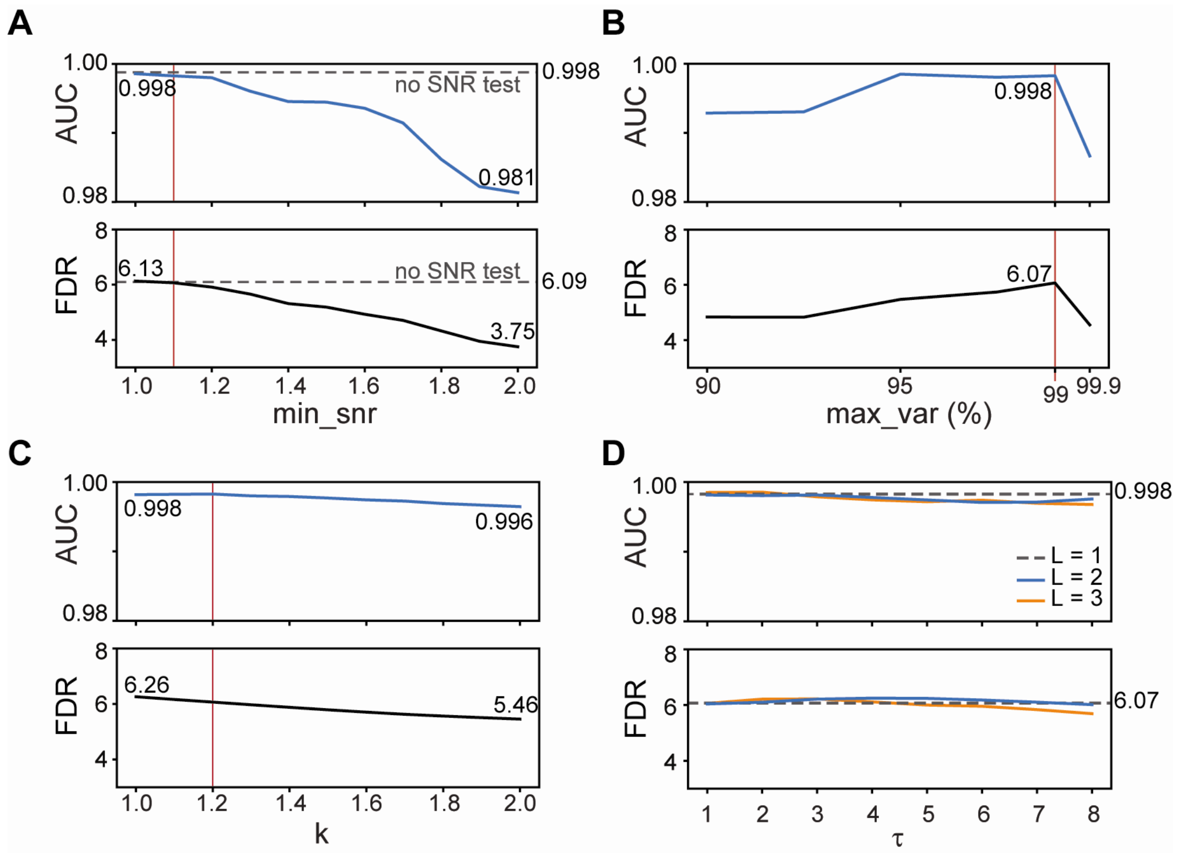


**Figure S2. Dependence of PCI$^{ST}$ on features of the TMS evoked potential.** PCI$^{ST}$ calculated on TMS evoked potentials recorded from conscious (wakefulness) and unconscious (NREM sleep/anesthesia) healthy individuals using different sampling frequencies, number of EEG channels and number of trials. (A) Influence of the signal's sampling frequency on PCI$^{ST}$ obtained by downsampling the TEP signals. Shown are the boxplots (boxplots) with median and quartiles for conscious (red box) and unconscious (grey box) conditions, and corresponding classification power metrics for sampling frequencies varying from 200Hz to 700Hz (top) (AUC: area under the ROC curve; FDR: Fisher's Discriminant Ratio, calculated as the square of the difference between the means of the distributions divided by the sum of their variances). (B) Dependence of PCI$^{ST}$ on the number of trials ($n_t$) used to obtain the TMS evoked potential (TEP). For every session, surrogates TEP's were generated by randomly selecting $n_t$ trials. The boxplots with median and quartiles (bottom) of the mean PCI$^{ST}$ values across 25 surrogates for conscious (red box) and unconscious (grey box) conditions are displayed with corresponding FDR and AUC (top) for $n_t$ ranging from 50 to 200 trials. Red asterisks indicate significant comparisons (univariate ANOVA for repeated measurements followed by Bonferroni post hoc test). (C) Correlation between PCI$^{ST}$ values for conscious (red) and unconscious (grey) conditions, with respective AUC and FDR values for the resulting distributions, calculated using the original hd-EEG system (horizontal axis) and two simpler EEG setups: a standard 10-20 EEG system (vertical axis, top panel) and a setup with 8 EEG channels (vertical axis, bottom panel).

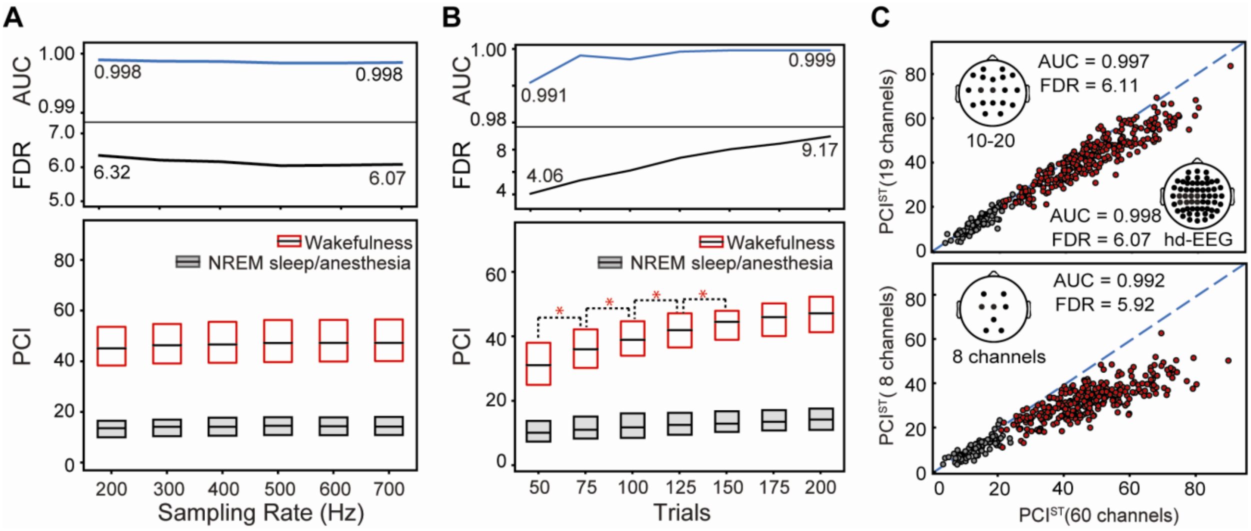


**Figure S3. Dependence of PCI$^{ST}$ on dimensionality reduction and state transition quantification parameters.** PCI$^{ST}$ calculated on TMS/EEG evoked potentials recorded from the benchmark population during wakefulness and NREM sleep/anesthesia for different varying parameters. Shown is the area under the curve (AUC) of the ROC curve and the Fisher's Discriminant Ratio (FDR) – calculated as the square of the difference between the means of the distributions divided by the sum of their variances – of conscious versus unconscious distributions. While a given parameter varied the remaining parameters were held constant at $k$ = 1.2, minimum SNR (min_snr) = 1.1, and maximum explained variance (max_var) = 99, and no time-delay embedding. Optimal parameters used for all calculations in the article are shown as the vertical red lines. (A) Influence on PCI$^{ST}$ of the signal-to-noise ratio test used in rejecting principal components with SNR < min_snr, where min_snr was varied from 1 to 2; the grey dotted line represents the case where no SNR test is performed. (B) Dependence of PCI$^{ST}$ on the total explained variance of the principal components calculated with SVD. Principal components where selected in order to add up to at least max_var% of the variance present in the response; max_var was varied from 90% to 99.9%. (C) Dependence of PCI$^{ST}$ on the noise control parameter k, varied from 1.0 to 2.0. (C) Effect of performing time-delay embedding to the signal before calculating PCI$^{ST}$ for embedding dimension L=2 (orange line) and L=3 (blue line), and delay τ varying from 1 to 8 time samples (1.4 ms to 11.0 ms), compared to using signal with no embedding (L=1, dotted grey line).

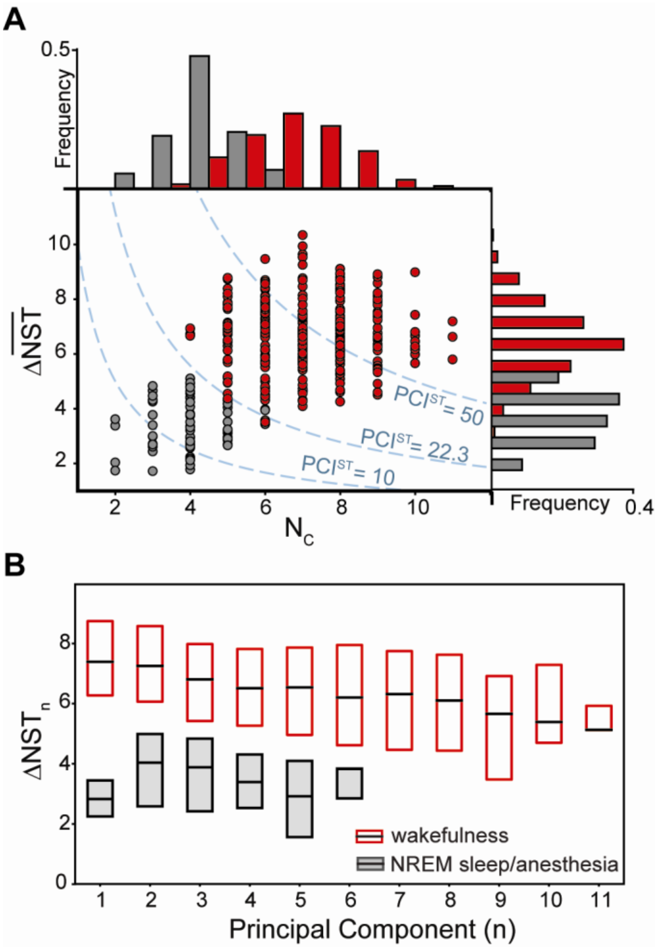


**Figure S4: Spatiotemporal complexity is captured by PCI$^{ST}$ through the number of selected components and the amount of state transitions for each component.** (A) In the center, the relationship between the number of principal components after dimensionality reduction ($N_C$) and the average number of state transitions across components ($\overline{\Delta \mathrm{NST}}$) for all sessions in the benchmark recorded during consciousness (red) and unconsciousness (grey). The former quantity can be regarded as a measure of the *spatial differentiation* of the brain's response to the perturbation, while the latter corresponds to the average *temporal complexity* present in the individual principal components as measured by the quantification of state transitions. The dotted blue lines correspond to the product of both quantities for different PCI$^{ST}$ values, since $\mathrm{PCI}^{\mathrm{ST}} = N_C \times \overline{\Delta \mathrm{NST}}$. On the right and on the top are the histograms of the average NST and of the number of selected principal components, respectively. (B) Relationship between selected principal component ordered by decreasing mean field power and the corresponding amount of state transitions for each component ($\Delta \mathrm{NST}_n$) (see Methods). Shown are the median and quartiles of $\Delta \mathrm{NST}_n$ values for conscious (red) and unconscious (grey) conditions for the $n$-th principal component, ranging from first to eleventh (as $N_C < 12$ for all sessions in the benchmark dataset).

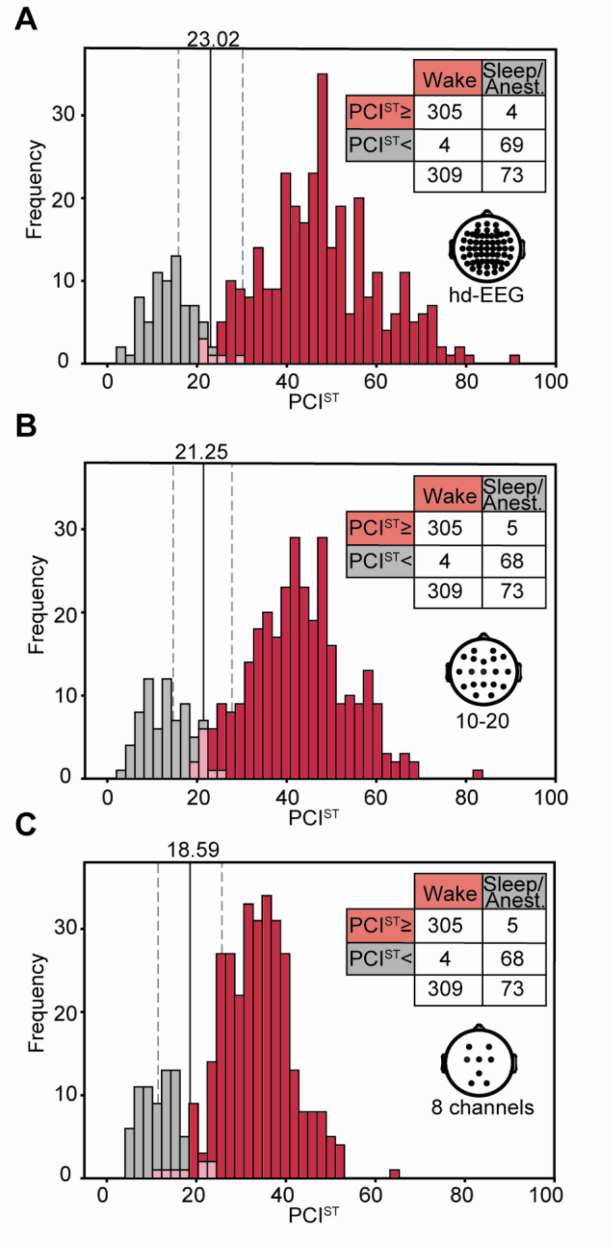


**Figure S5. Support vector machine (SVM) threshold calculated on benchmark population for different EEG setups.** In order to classify the brain-injured patients from the DOC population, a statistical threshold was extracted from the distribution of $PCI^{ST}$ values of all TMS/EEG session from the benchmark. This threshold was obtained using support vector machine, a technique that looks for a separator of the labeled data that maximizes the distance from the nearest data points. We employed the LinearSVC() function from the sklearn.svm module of the Python scikit-learn library using maximum number of iterations = 10000 and standard values for the remaining parameters (penalty parameter C=1). Displayed are the distributions of $PCI^{ST}$ for all TMS/EEG sessions in the benchmark dataset (NREM sleep/anesthesia sessions in grey, wakefulness session in red; overlaps are shown in pink) for EEG setups of 60 (A), 19 (B) and 8 (C) channels. The resulting SVM thresholds (numbers in the panels and grey solid lines) are shown together with their respective SVM margins (dotted grey lines).

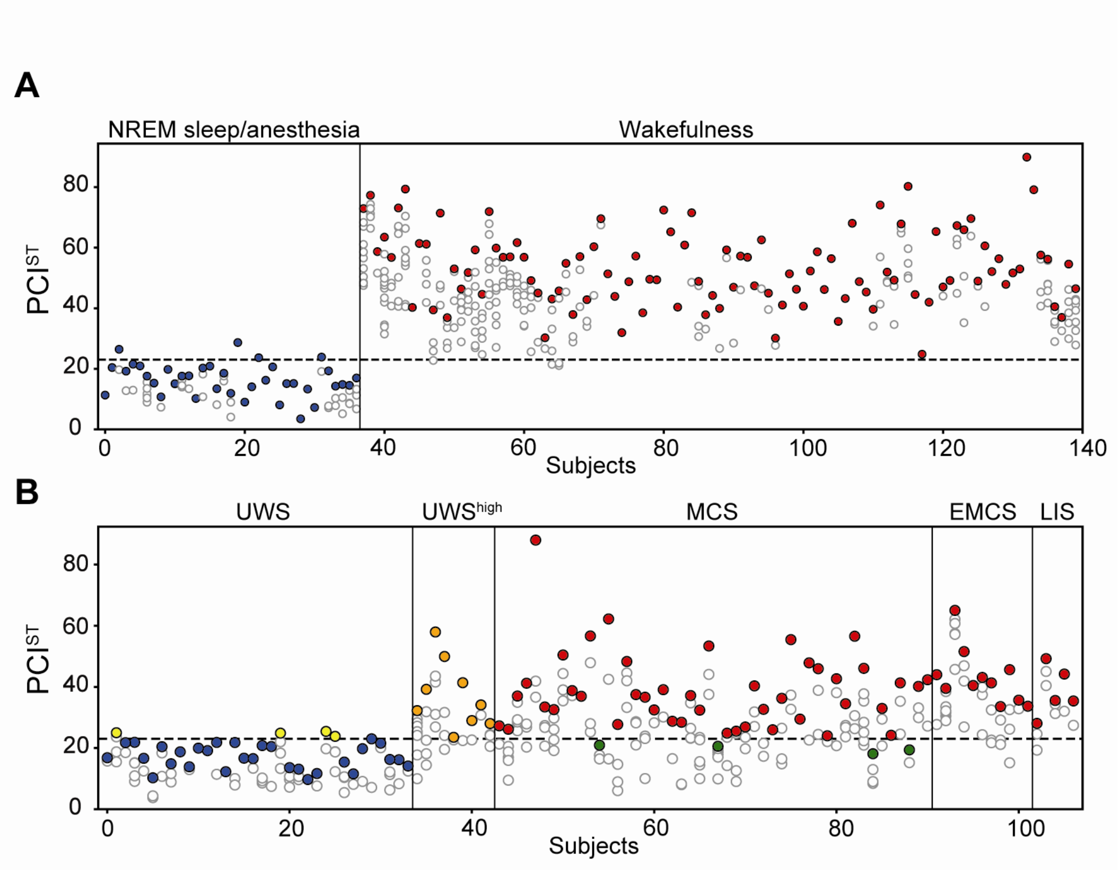


**Figure S6. PCI$^{ST}$'s classification of the benchmark and DOC population according to SVM threshold.** Scatter plots show PCI$^{ST}$ values for each TMS/hd-EEG sessions arranged by condition and subject. Each dot corresponds to the PCI$^{ST}$ value for a given TMS/hd-EEG session, and dots in the same column are sessions recorded from the same subject. The subject's maximum PCI$^{ST}$ value is represented with a filled dot colored according to the condition and to its position in respect to the threshold extracted from the benchmark (see S4 Fig). (A) PCI$^{ST}$ values for all healthy subjects in the benchmark population, which includes sessions recorded during alert wakefulness, NREM sleep, and anesthesia sedation (midazolam, xenon and propofol). (B) PCI$^{ST}$ results for all sessions from the DOC populations, comprised of patients with unresponsive wakefulness syndrome (UWS), minimally conscious state (MCS), emerging minimally conscious state (EMCS) and locked-in syndrome (LIS). In particular, UWS patients which presented high values of complexity as previously measured by PCI$^{LZ}$ are shown separately as UWS$^{high}$ .

**Table S1:** Anatomical regions of individual SPES contacts in correspondence to Figure 5.

| Subject | Contact Number | Anatomical Region |
|---|---|---|
| #1 | 1 | Superior frontal gyrus |
| | 2 | Lateral occipito-temporal gyrus (fusiform gyrus) |
| | 3 | Central sulcus |
| | 4 | Superior frontal gyrus |
| | 5 | Inferior frontal sulcus |
| | 6 | Middle occipital gyrus |
| #2 | 1 | Middle-anterior part of the cingulate gyrus and sulcus |
| | 2 | Subparietal sulcus |
| | 3 | Precuneus |
| | 4 | Posterior-dorsal part of the cingulate gyrus |
| | 5 | Sulcus intermedius primus (of Jensen) |
| #3 | 1 | Subcentral gyrus and sulci |
| | 2 | Inferior part of the precentral sulcus |
| | 3 | Inferior part of the precentral sulcus |
| | 4 | Middle-anterior part of the cingulate gyrus and sulcus |
| | 5 | Middle-anterior part of the cingulate gyrus and sulcus |
| | 6 | Hippocampus |
| | 7 | Hippocampus |
| #4 | 1 | Transverse temporal sulcus |
| | 2 | Posterior ramus of the lateral sulcus |
| | 3 | Intraparietal sulcus and transverse parietal sulci |
| | 4 | Middle-posterior part of the cingulate gyrus and sulcus |
| | 5 | Intraparietal sulcus and transverse parietal sulci |
| | 6 | Hippocampus |
| | 7 | Superior occipital sulcus and transverse occipital sulcus |
| | 8 | Posterior ramus of the lateral sulcus |
| #5 | 1 | Superior frontal gyrus |
| | 2 | Superior frontal sulcus |
| | 3 | Middle frontal gyrus |
| | 4 | Parahippocampal part of the medial occipito-temporal gyrus |
| | 5 | Middle-anterior part of the cingulate gyrus and sulcus |
| #6 | 1 | Posterior-dorsal part of the cingulate gyrus |
| | 2 | Superior frontal sulcus |
| | 3 | Superior frontal gyrus |
| | 4 | Middle frontal gyrus |
| #7 | 1 | Inferior frontal sulcus |
| | 2 | Opercular part of the inferior frontal gyrus |
| #8 | 1 | Middle-anterior part of the cingulate gyrus and sulcus |
| #9 | 1 | Superior frontal sulcus |
| | 2 | Orbital sulci (H-shaped) |
| | 4 | Superior segment of the circular sulcus of the insula |
| | 3 | Middle-anterior part of the cingulate gyrus and sulcus |
